# Improvement of bone regeneration using fibrin biopolymer combined with differentiated stem cells

**DOI:** 10.1101/608166

**Authors:** Camila Fernanda Zorzella Creste, Patrícia Rodrigues Orsi, Fernanda Cruz Landim-Alvarenga, Luis Antônio Justulin, Marjorie de Assis Golim, Benedito Barraviera, Rui Seabra Ferreira

## Abstract

Bone tissue repair remains a challenge on tissue engineering. New approaches are highly expected to regenerate fractures, bone infections, cancers and congenital skeletal abnormalities. Lately, osteoconductive biomaterials have been used with osteoprogenitor cellsas bone substitutes to accelerate bone formation. Fibrin scaffold serves as a provisional platform promoting cell migration and proliferation, angiogenesis, connective tissue formation and growth factors stimulation. When combined with mesenchymal stem cells (MSCs) maintain cell viability that exerts an immunomodulatory effect by modifying inflammatory environment through expression of pro and anti-inflammatory cytokines. We evaluated a unique heterologous fibrin biopolymer as scaffold to MSCs on bone regeneration of rat femurs. A critical-size bone defect was made in the femur and treated with fibrin biopolymer(FBP); FBP + MSC; and FBP + MSC differentiated in bone lineage (MSC-D). Bone repair was analyzed 03, 21 and 42 days later by radiographic, histological and scanning electron microscopy (SEM) imaging. The FBP+MSC-D association was the most effective treatment, since newly formed bone was more abundant and early matured in just 21 days. Our results demonstrate that FBP isolated was able to promote bone repair although cells play a crucial role on the type and quantity of bone tissue formed. We have not observed surgical site infection, inflammatory response, fractures or loss of function related with FBP. Thus, this approach can be safely expanded for clinical trials as an effort to overcome current method limitations and improve overall bone regeneration process.

## 2. BACKGROUND

Bone tissue repair is frequently necessary after skeletal diseases, congenital abnormalities, infections, trauma and surgical procedures after hematological, breast and ovary cancers. Fractures with bone loss often require grafts or implants. Autologous and allogeneic grafts represent about 90% of bone tissue transplants while inorganic matrices represents the other 10%^1,2^. Ideal implants must act as scaffold for bone regeneration with host tissue integration.

Main function of scaffolds is to offer structure and support for migration and specialization of different cells involved in healing. This structure should allow cell adhesion, attachment, differentiation, proliferation and biological function for repair of the injured tissue^3^. Synthetic osteoconductive implants such as hydroxyapatite and calcium triphosphate have porous structures that promotes bone growth, however, the absence of an osteoinductive potential is still a limitation^4^.

Mesenchymal stem cells (MSC) are used in tissue engineering^5,6,7^ as an excellent alternative for bone repair since they are able to differentiate in osteoblastsas also in condrocytes, myocytes, adipocytes and fibroblasts^8^. MSC applied in tissue repair has evolved progressively to improve or even substitute the healing capacity of bone tissue in partial or complete failure of the repair mechanism^9,10^.

Combination of live cells with synthetic or natural scaffolds has been used to produce live tridimensional tissues that are functional, structural and mechanically identical to the original^11, 12,13^. Different compounds have been used as scaffolds for MSC^14^ and can be classified as synthetic (i.e., hydroxyapatite and calcium triphosphate)^15^ or biological as fibrin biopolymers^16, 17^.

Commercially available fibrin biopolymers are used in different surgical fields as hemostatic agents, healing promoters, cavity sealers anddrug delivery in surgical sites^18,19^. Fibrin biopolymers have showed *in vitro* similar structure and mechanical properties to those of the fibrin clot *in vivo*^20, 21^.

Biocompatibility, biodegradability and the capacity to interact with MSC suggest that fibrin biopolymers are important vehicles for cell transplantation^20,21,22^. However, they are derived from human thrombin and fibrinogen which has a risk of infectious disease transmission and limited use due to possible lack of the main components^20,23, 24, 25^.

A new fibrin biopolymer (FBP) constituted of two animal derived compounds instead of human blood has been used in experimental biomedical applications^24,26, 27, 28, 29, 30, 31^, such as nervous tissue^32,33^ and bone^34^ repair as also on the treatment of chronic venous ulcers in human patients^30,33^.

This FBPenabled*in vitro* MSC adhesion and growth and had no negative effect on osteogenic, adipogenic and chondrogenic differentiation. It also showed that FBP surface enhanced specific markers of cell viability^16^.

Although many associations of scaffolds and MSC are being studied for bone defect healing^35, 36, 37,38^ there are still challenges to be faced. Aiming to overcome current method limitations we evaluated the effect of this new FBP with MSC and osteogê nica differentiated MSC on the treatment of critical-size defects in rats.

FBP+MSC-D association has showed most effective treatment, since newly formed bone was more abundant and early matured in just 21 days.

## 3. MATERIALS AND METHODS

This study was approved by Experimental Ethics Committee of Botucatu Medical School, Sã o Paulo State University, Brazil (n° 968-12).

### Fibrin Biopolymer (FBP)

The FBP was kindly provided by Center for the Study of Venoms and Venomous Animals (CEVAP), UNESP, Botucatu, SP, Brazil. Components were distributed in three vials containing thrombin-like enzyme, animal cryoprecipitate and diluent and were kept frozen at -20°C until use^34,39, 40, 41, 42^. At time of surgery contents were immediately mixed according to the manufacturer’s package insert.

### Cell isolation and culture

Cell isolation and culture was performed according to Orsi *et al*., 2017^34^. Twelve Wistar rats 10days-oldwere euthanized with halothane overdose (MAC> 5%) and used as bone marrow donors. Marrow cells were obtained from the femur by insertion of needle syringe into the bone cavity and then washing with Dulbecco’s modified Eagle medium (DMEM) (Gibco Laboratories, Grand Island, NY, USA).

Pool of material was centrifuged at 2000rpm for 10min after bone marrow harvest and re-suspended in DMEM (Gibco Laboratories, Grand Island, NY, USA) supplemented with 20% fetal bovine serum (Sigma-Aldrich, St. Louis, MO, USA), 100 μg/mL of penicillin/streptomycin solution (Gibco Laboratories) and 3 μg/mL of amphotericin B (Gibco Laboratories, Grand Island, NY, USA).

Cells were seeded in 75 cm^2^ culture flasks and placed in a 5% CO_2_incubator at 37.5°C. Culture medium was changed every 03 days and cell growth and adherence were monitored by inverted microscopy. Cells were subcultured when reached 80% confluence. All experiments were performed with MSC at passage 3 (P3). To passages, culture medium was discarded, cells were washed with 02mL of PBS followed by addition of trypsin solution (Gibco, Grand Island, NY, USA) and 05min incubation at 37.5°C. Thus were centrifuged for 10min at 2000rpmand re-suspended in culture media. Cells were counted and 1×10^6^cells/dose were used in association with FBP for the treatment of the bone defect throughout the experiment.

Cells were characterized by flow cytometry (FACSCalibur; BD Pharmingen, San Diego, CA, USA) using monoclonal antibodies for specific positive and negative markers (Table 1)^13, 14, 43, 44^. Assays were performed using 2×10^5^ cells and data were analyzed using the CellQuest Pro^®^ software after acquisition of 20,000 events. Functional characterization was also performed as cells were differentiated in osteogenic, chondrogenic and adipogenic lineages after the third passage^22, 35, 45^.

**Table 1:** Surface markers for MSC characterization.

| Negative Markers |  |
| --- | --- |
| RT1 | anti-RT1-Aw2-FITC, clone MRC OX-18; Abcam, Cambridge, MA, USA |
| CD34 | anti-CD34-PE, clone ICO-115; Abcam, Cambridge, MA, USA |
| CD11b | anti-CD11b-PE, clone ED8; Abcam, Cambridge, MA, USA |
| CD45 | anti-CD45-FITC, clone MRC OX-1; Abcam, Cambridge, MA, USA |
| MHCII | anti-rat MHC CLASS II RT1D-PE, clone MRC OX-17; Abcam, Cambridge, MA, USA |
| Positive Markers |  |
| CD73 | purified mouse anti-rat CD73; clone 5F/B9, BD Pharmingen, San Diego, CA, USA |
| CD90 | anti-CD90/Thy1-FITC, clone FITC.MRC OX-7; Abcam, Cambridge, MA, USA |
| CD44 | anti-CD44-PE, clone OX-50; Abcam, Cambridge, MA, USA |
| ICAM-I | anti-ICAM-I-FITC, clone 1A29; Abcam, Cambridge, MA, USA |

### Osteogenic differentiation of MSC

After cell culture had reached 70% confluence, culture medium was replaced by StemPro® Osteogenesis Differentiation Kit medium (Gibco by Life Technologies A10072-01, Carlsbad, CA, USA), composed of 73% Osteocyte/Chondrocyte Differentiation Basal Medium (Gibco by Life Technologies A10069-01, Carlsbad, CA, USA), 5% Osteogenesis Supplement (Gibco by Life Technologies A10066-01, Carlsbad, CA, USA), 1% penicillin/streptomycin, 1% amphotericin B, and 20% fetal bovine serum (Sigma-Aldrich, St. Louis, MO, USA). The differentiation medium was replaced every 03 days for 12 days.

Then, cells were fixed in ice-cold 70% ethanol, washed in distilled water and stained in 02 mL of Alizarin red (Invitrogen Life Science Technologies, Carlsbad, CA, USA) for 30min at room temperature. After the dye was removed, cells were washed four times in distilled water and observed in an inverted light microscope^17,46^.

### Animals and surgical protocols

Twenty-seven male Wistar rats (*Rattus norvegicus*) weighing 200 to 250g were kept at 21 ± 2 °C under a 12h light/dark cycle and allowed access to food and water *ad libitum*. Surgeries were carried out under sterile conditions.

Animals were weighed and anesthetized with ketamine solution (1mL/Kg) and xylazine hydrochloride (0.25mL/Kg) intraperitoneally. Cross section of the thigh through the upper- and middle-third of the femur allowed a critical defect of 05 mm to be performed on the distal epiphysis of the right femur with a low rotation drill (BELTEC^®^) under constant irrigation of 0.9% sterile saline to prevent overheating^47^. Postoperative analgesia with intramuscular flunixin-meglumine (1mg/Kg) was performed every 24h for three days.

Animals were distributed in three experimental groups of 09 animals each: control group (CG) were animals were treated with FBP only; Group 1 (G1) were animals were treated with FBP + MSC; and Group 2 (G2) that received FBP + MSC-D). Cells were mixed in 100 μLFBP immediately before injection at 1×10^6^ cells/dose for G1 and G2.

### Radiographic evaluation

Radiographic imaging of the rat femurs was conducted at 3^rd^, 21^th^ and 42^th^ days using a digital GE model E7843X system (GE Healthcare, Chicago, IL, USA).

### Histological analysis

Femurs were removed and fixed in 10% buffered formalin for 24h at 4°C, and were decalcified with 10% neutralized EDTA (Sigma^®^) for 04 weeks; then dehydrated with an ascending series of ethanol concentrations, cleared in xylene, and embedded in Paraplast^®^ (Sigma^®^). Histological sections (06μm) were stained with hematoxylin–eosin (H& E) for general morphological analysis or picrosirius for collagen fibers (type I and type III) quantification and stereological analysis^46^. The color displayed under polarizing microscopy, is a result of fiber thickness, as well as the arrangement and packing of the collagen molecules. Normal tightly packed thick collagen fibers have polarization colors in the red spectrum while thin or unpacked fibers have green birefringes^48^. Sections were observed under normal and polarized light, and digitalized images were analyzed using Leica Q-win^®^ software (version 3.0) to calculate mean collagen fiber area.

### Scanning electron microscopy (SEM)

SEM analyses were performed using a Quanta 200 electron microscope (FEI Company, Hillsboro, OR, USA). Bone samples were fixed in 2.5% glutaraldehyde in 0.1 M PBS pH 7.3 for 04h. The samples were then removed and washed three times for 05min in distilled water. Subsequently, samples were immersed for approximately 40min in 0.5% osmium tetroxideand washed three times in distilled water; dehydrated in increasing concentrations of ethanol (7.5% to 100%); dried in a critical point apparatus with liquid carbon dioxide, mounted on appropriate chucks, metallized and gold-coated^36^.

## 4. RESULTS

### MSC expansion and characterization

MSC exhibited fibroblastoid morphology (Fig. 01). Cells remained in primary culture until reached 80% confluence after approximately 07 days; then subcultured up to the third passage for use. Flow cytometry showed that 97.57%, 98.49%, 84.47% and 91,70% of the cells expressed positive markers ICAM-I, CD90, CD73, and CD44, respectively (Fig 02). Negative markers MHC II, CD34, CD45, RT-1, and CD11b were expressed respectively by 1.45%, 1.32%, 2.39%, 1.80%, and 1.74% of cells (Fig. 03). These results demonstrates that cultured cells exhibited the characteristic phenotype of MSC.

**Fig. 01.**
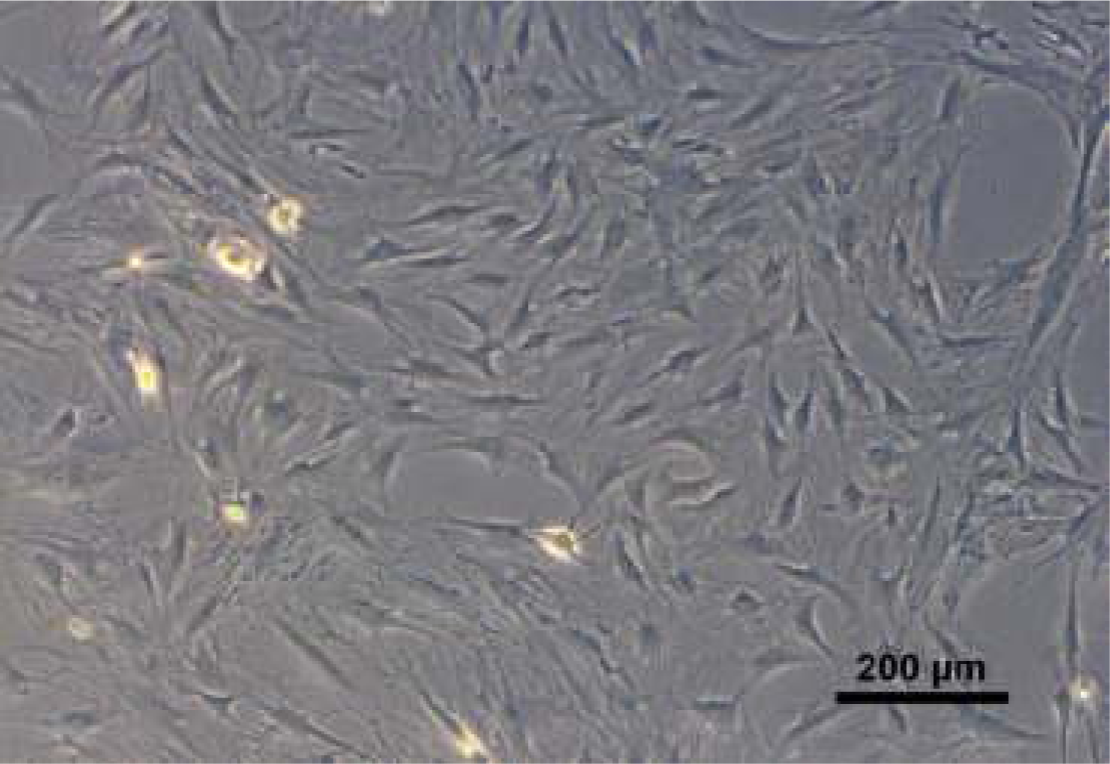
Cultivated MSC presented expected fibroblastoid (fusiform) shape.

**Fig. 02.**
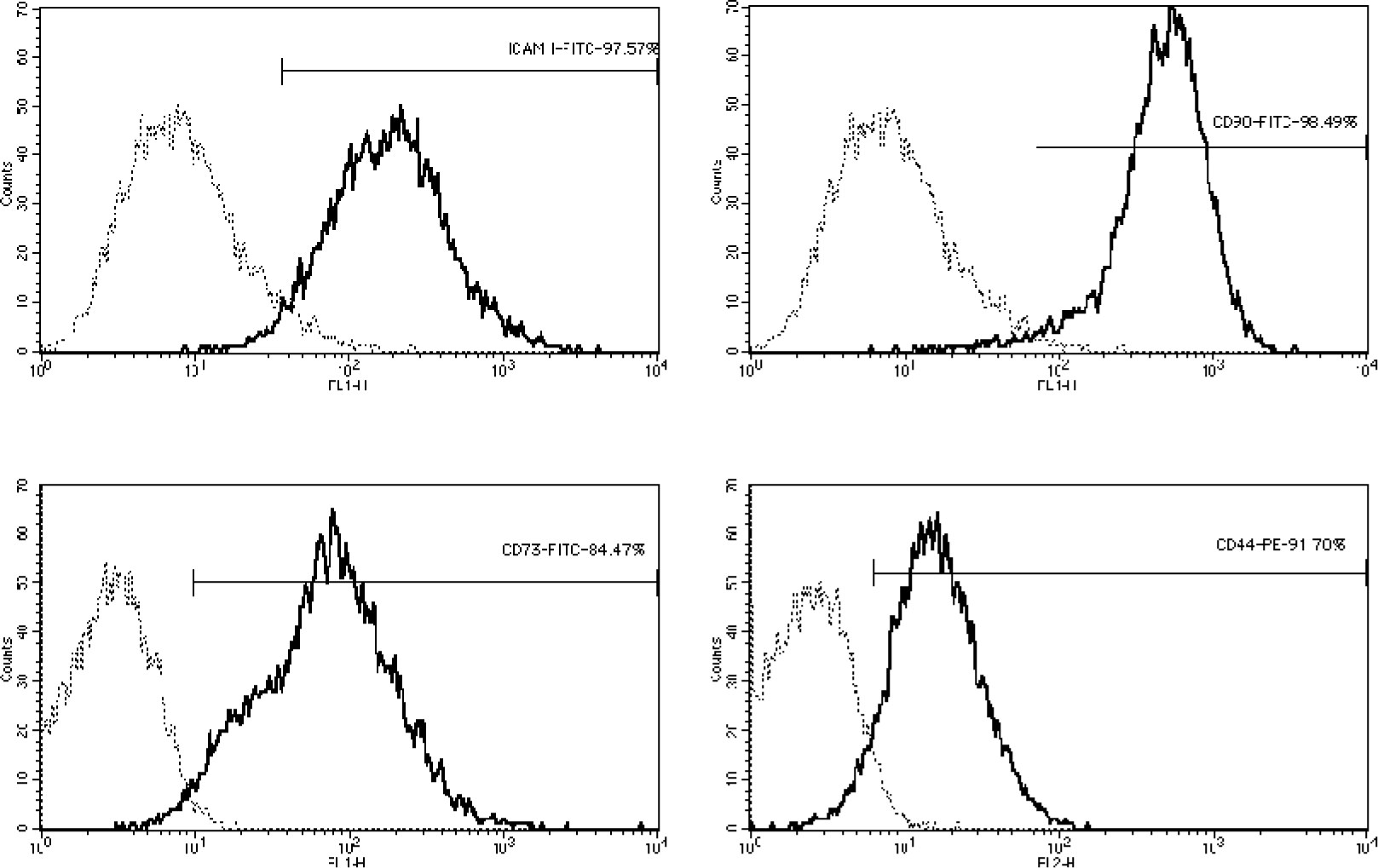
Positive markers on flow cytometry analysis. Upper left – ICAM-I; Upper right – CD90; Lower left - CD73; Lower right – CD44.

**Fig. 03.**
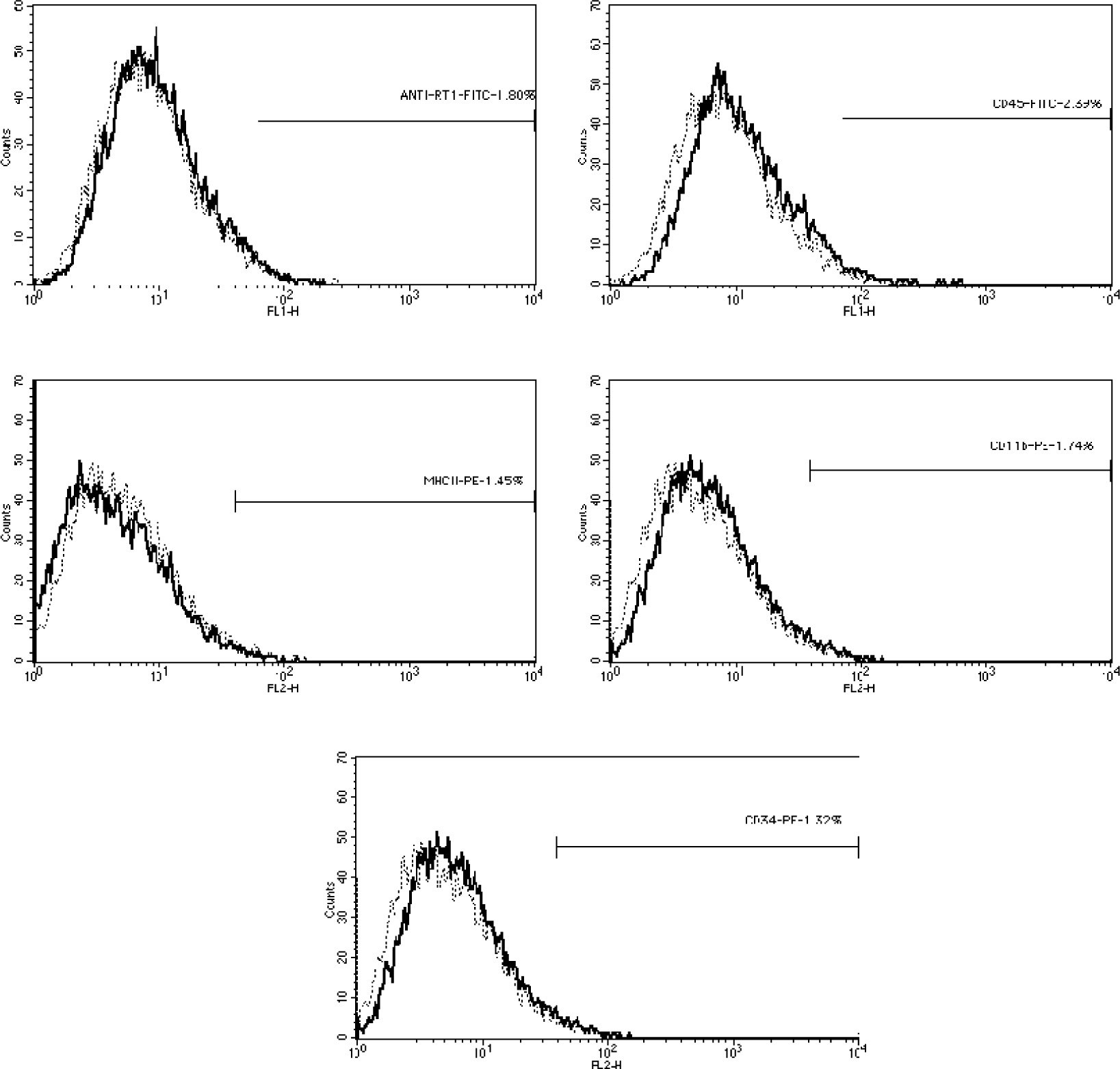
Negative markers on flow cytometry analysis. Upper left – anti-RT1; Upper right – CD45; Lower left – MHC II; Lower right – CD11b; Center – CD34.

### MSCs osteogenic differentiation

Figure 04 shows calcium deposits observed in MSC cultures after 12 days of incubation in specific differentiation media. Mineral deposits were detected by presence of red staining on the extracellular medium, thus confirming the MSC osteogenic differentiation.

**Fig. 04.**
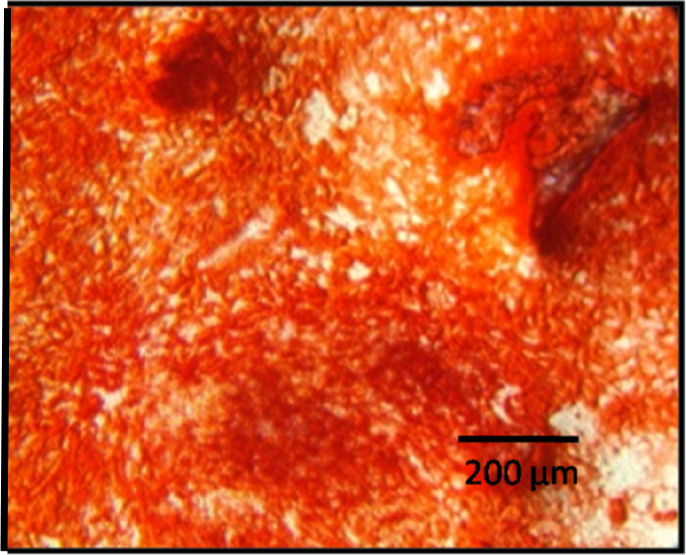
Calcium deposits stained red in MSC cultures after 12 days of differentiation.

**Fig. 05.**
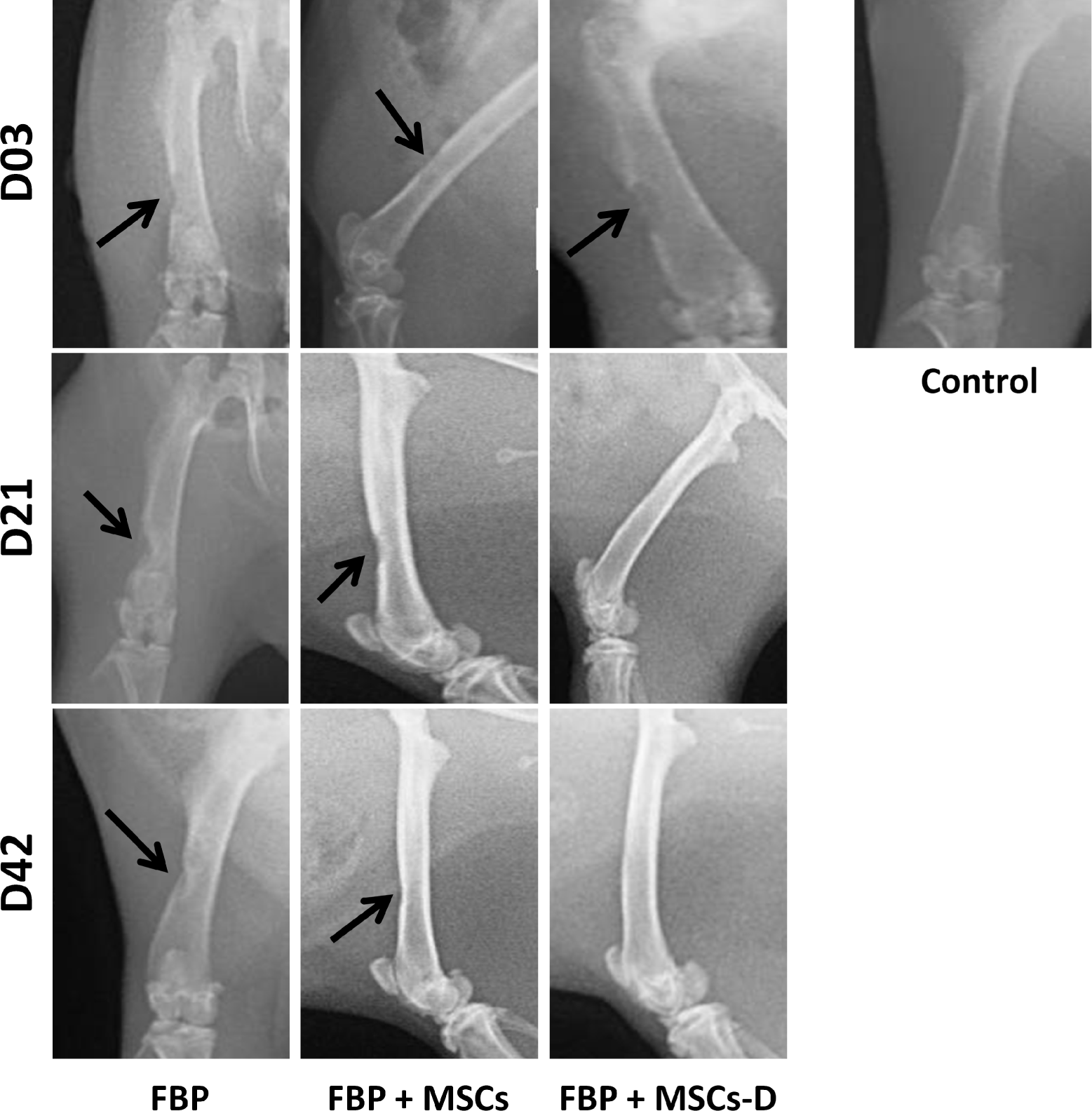
Radiographic analysis of injury site at 03 different time points.

### Radiographic evaluation

Radiographic analyses showed that defects were evident in all groups 03 days after surgical procedure. At the 21^st^ day, FBP+MSC-D treated group presented efficient healing as the defect was almost completely filled. At day 42, FBP + MSC-D group showed total bone healing and it was possible to observe improve of repair on FBP + MSC treated group.

### Scanning electron microscopy (SEM)

Scanning electron microscopy imaging evidenced the bone structure at injury site. On the 3^rd^ day after surgery defect was evident in all groups. Group treated with FBP+MSC-D showed markedly higher injury repair when compared to the other two groups at day 21. After 42days it was possible to observe bone tissue deposits in all treated groups. However, in groups FBP and FBP + MSC the defect has not been completely repaired as could be observed on group FBP+MSC-D (Figure 06).

**Fig. 06.**
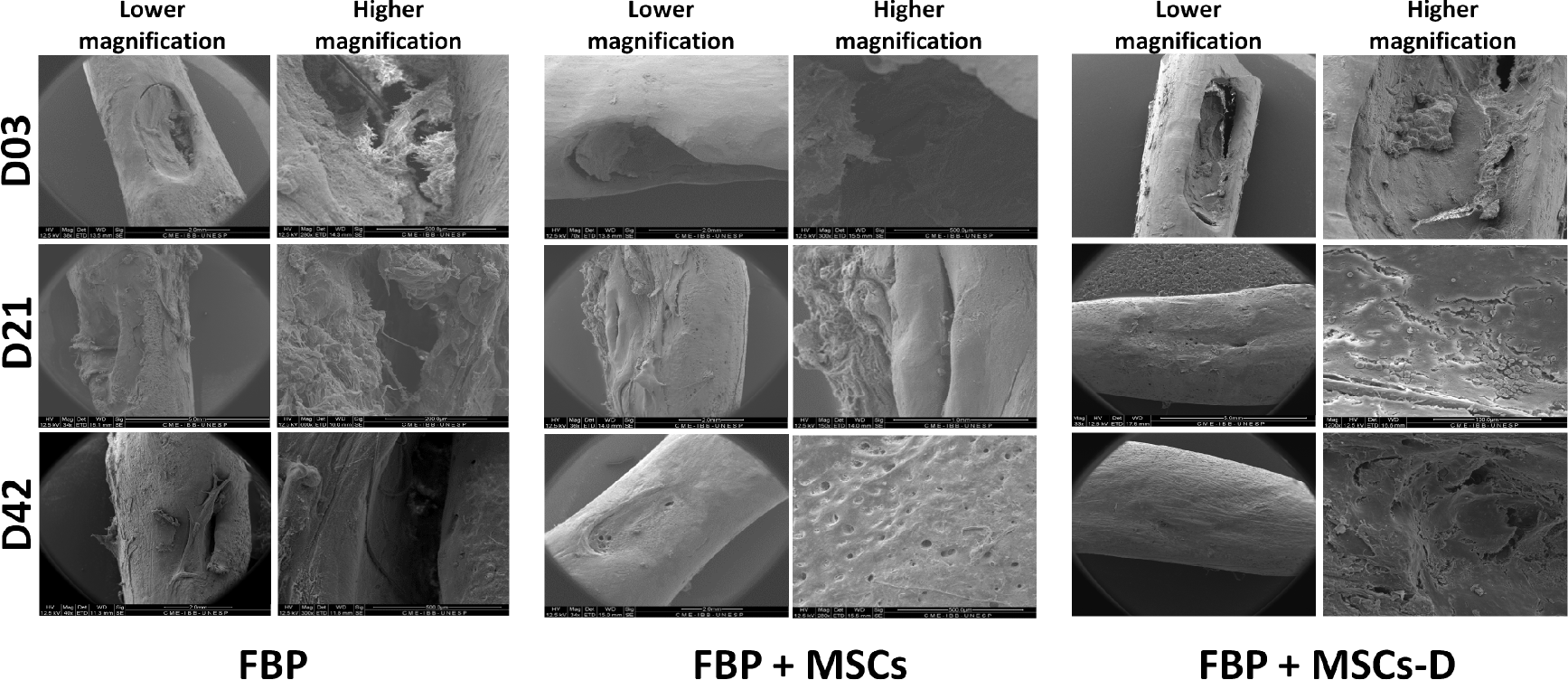
SEM imaging of injury site of the studied groups at 03 different time points.

### Histological analysis

H& E stained material are demonstrated in Figure 07. A progressive bone matrix deposition was observed during the experimental period. Presence of a fibrilar material, similar to FBP structure, insidethe defect 03 days after surgery on the group treated only with FBPevidences that it has adhered to injury site. Bone fragments probably from the surgical procedure were also observed adhered to the fibers. There was a significa t increase in cellullarity associated to the biomaterial. On the group treated with FBP + MSC the presence of newly formed trabecullar bone on the defect margins was evident after 21 days as well as in the FBP + MSC-D treated group. At day 42, from histological perspective, all defects were partially repaired, although in the FBP + MSC-D treated group newly formed bone was more abundant and its structure more similar to normal mature bone tissue.

**Fig. 07.**
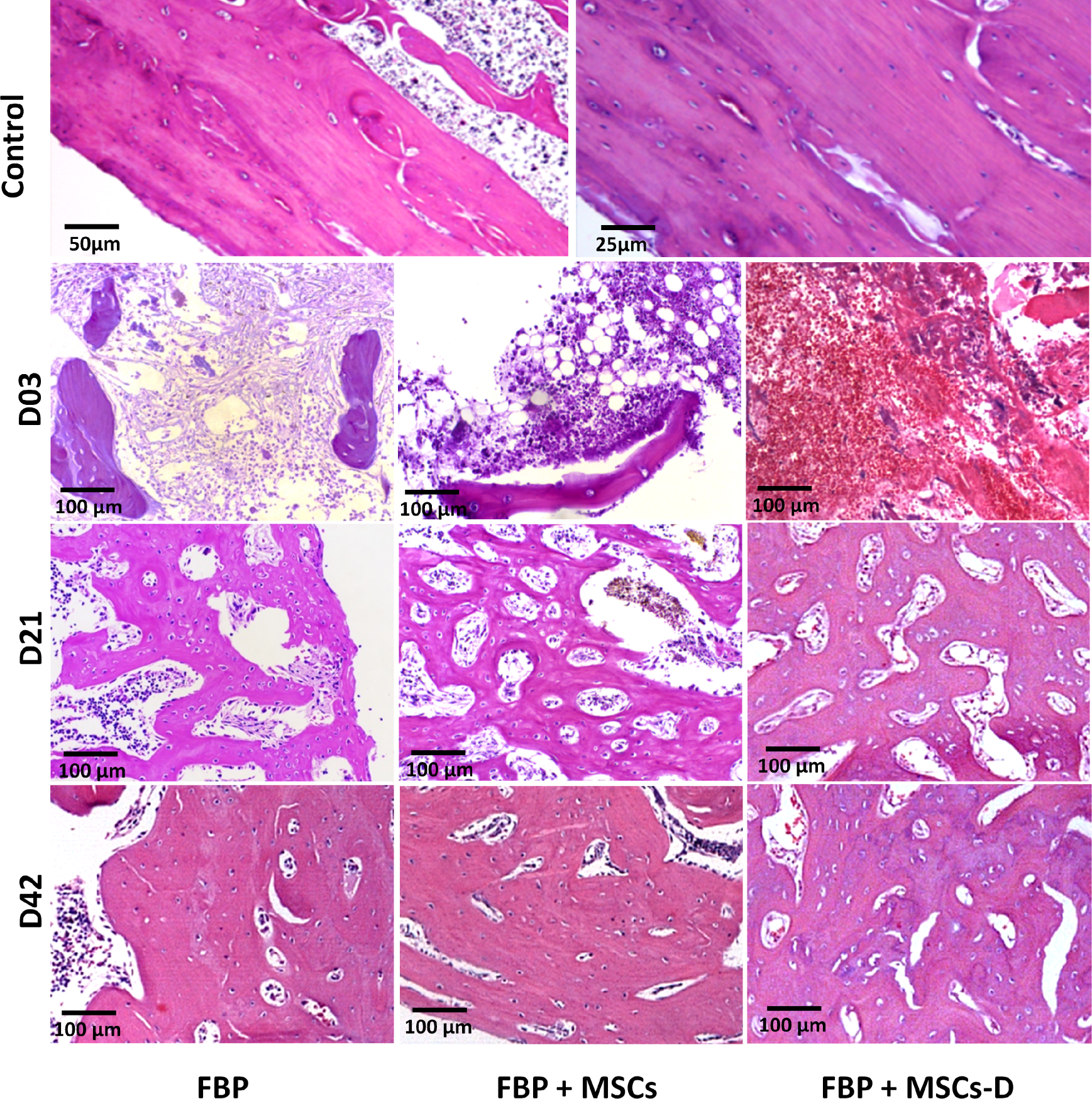
Histological analysis of bone regeneration tissue stained with H& E.

Figure 08 shows collagen fibers formation through Picrosirius staining under polarized light. Yellow-reddish staining represents mature thick fibers, as demonstrated by the control group, while green staining shows recent synthesized and immature fibers. On the first 03 days there was no evidence of collagen formation in all groups. After 21 days, there were observed thin immature green fibers within thick yellow and red mature collagen showing an increase in collagen synthesis in all three groups. Besides, there were also a high number of cells adhered to the scaffold claiming that MSC injected with the FBP remained at injury site and have differentiated for matrix synthesis. Our results shows that even after 21 days cells were associated with the fibrin structure strengthening the use of FBP as a scaffold for cell delivery. The same collagen synthesis pattern was observed in all groups at day 42, thus it was higher on group FBP + MSC-D.

**Fig. 08.**
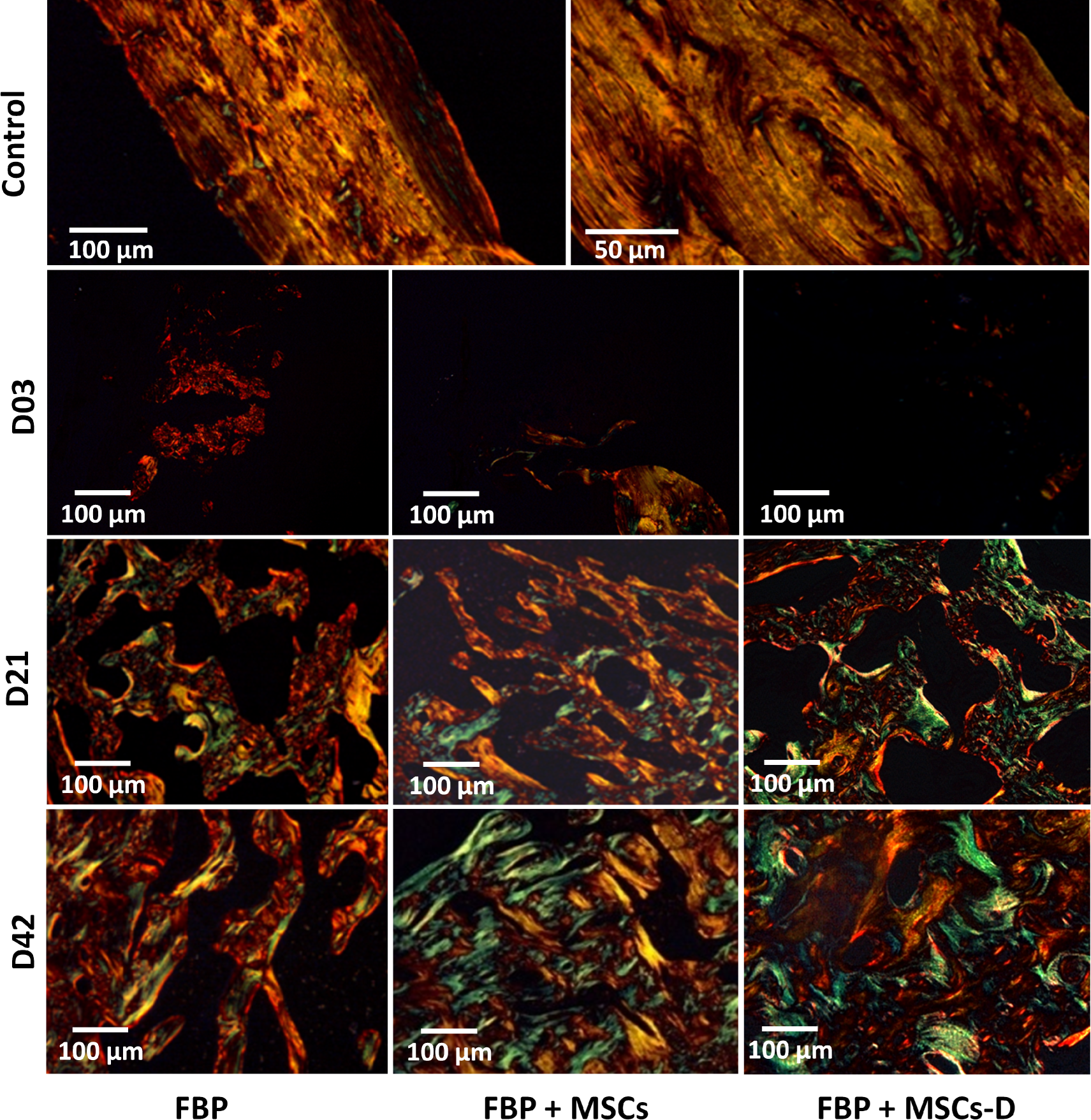
Picrosirius staining under polarized light showing collagen formation in injury filling.

## 5. DISCUSSION

Although autologous bone graft remains the gold standard for healing large bone defects, grafting procedure complexity increases due to donor site morbidity, increased risk of infection and poor ability to fill complex defects^49^, besides the feasibility to obtain material in adequate quantity and quality. However, the autograft has its limitations, including donor-site morbidity and supply limitations, hindering this as an option for bone repair^50^.

Scaffolds to bone tissue repair must induce bone formation and provide a suitable microenvironment for growth of bone cells exhibiting osteoconductivity, osteogenicity and osteoinductivity^50^. So, an ideal scaffold requires adequate porosity (at least 30%) and pore size (at least 100 μm in diameter) to facilitate cell migration, differentiation and ingrowth within the scaffold^51, 52, 50^.

Our results demonstrate *in vivo* the potential of fibrin biopolymer (BPF) to act as a scaffold for MSC in bone repair evidencing its biocompatibility. Association between BPF and MSC-D was able to promote total repair in critical size defect in rat femurs in almost half-time when compared to other studied treatments.

Commercially available fibrin biopolymers, also called fibrin sealants, consist of human fibrinogen and thrombin. The BPF used in this study is composed of a mixture of a serine protease with thrombin-like enzyme activity, purified from *Crotalus durissus terrificus* snake venom and buffaloes cryoprecipitate as a source of fibrinogen^53, 30^. Previous studies has shown no cytotoxicity becoming an excellent scaffold for MSC^16, 34, 54, 55^.

Previous studies have shown that BPF promotes chemotaxis for M2 macrophages which presents anti-inflammatory profile and promotes neoangiogenesis^31^. We did not observe signs of local inflammation proved by animals postoperative status with normal cicatrization and absence of flogistic signs of inflammation and surgical site infection such as erythema, local edema or exudates. Also, there were few leukocyte infiltrates that are characteristics of foreign body reactions evidencing BPF biocompatibility.

According to Ahmed *et al*.^56^, fibrin derivates used in tissue engineering are hidrogels, sealants and fibrin microspheres. Gasparotto *et al.*^16^ demonstrated *in vitro* interactions of the heterologous fibrin sealant (BPF) with MSC either in scanning electron microscopy (SEM) or in transmission electron microscopy (TEM). Authors concluded that BPF showed ideal plasticity and MSC homing without differentiation effects. Orsi *et al.*^34^, have evaluated the effect of BPF associated with both MSC and MSC-D on osteoporotic female rats and showed that the association promotes a higher bone formation compared to the control group after 14 days. They also have demonstrated that there was no cytotoxicity of BPF for MSC.

MSC characterization was effective either in microscopy or in flow citometry (FC). Cells presented expected fusiform shape in culture and FC panel chosen was adequate and agreed with other authors that stated MSC should present positive for CD73, CD90, CD105 e ICAM and negative for CD45, CD34, CD14 or CD11b, CD79 or CD19^57, 58, 59, 60^. Additionally, rat bone marrow derived MSC have differentiated in osteogenic lineage after 12 days on presence of specific differentiation media corroborating Vilquin & Rosset, 2006^9^.

BPF helped cicatricial evolution with total wound healing after observational period. Group treated with BPF and MSC-D highlighted from the others as it presented complete repair after 21 days. Xu *et al.*^47^ also evaluated a new scaffold composed by BG-COL-HYA-OS and MSC in rat femur regeneration and have observed a significant injury filling after 42 days.

BPF has also been used as a scaffold in the regeneration of other tissues. Cartarozzi *et al.*^32^ have investigated in rats the effectiveness of MSC associated with a fibrin scaffold for the regenerative process after peripheral nerve tubulization and showed that the association improves nerve regeneration by positively modulating the reactivity of Schwann cells^32^.

Spejo *et al.*^55^ have demonstrated that MSC therapy is neuroprotective and when associated with a BPF shifts the immune response to a proinflammatory profile. According to them, thet was possible because BPF kept EGFP-MSC at the glial scar region in the ventral funiculus after 28 days.

Radiographic analysis is an auxiliary measure for repair evaluation in bone lesions as it provides neither information about bone quality in new tissue nor it allows for a clear visualization of old-new bone interface^61^. Histological and SEM analysis confirmed radiographic findings and also complemented the information. Considering the three analysis allowed us to conclude that the association between BPF and MSC-D was able to promote total repair in critical size defect in rat femur and shortened bone repair compared to other evaluated treatments.

BPF presented as a highly effective scaffold for applications in bone lesions because it accelerated tissue regeneration. We believe that it was due to the use of mesenchymal stem cells pre differentiated in bone lineage associated with scaffold ability to keep cells viable on injury site without any adverse events.

## REFERENCES

1 Weir M.D., Xu H.H.K. Human bone marrow stem cell-encapsulating calcium Phosphate scaffolds for bone repair. ActaBiomater. 6, 4118–26 (2010).

2 Bressan E. Biopolymers for Hard and Soft Engineered Tissues: Application in Odontoiatric and Plastic Surgery Field. Polymers. 3, 509–526 (2011).

3 Stock U.A., Vacanti J.P. Tissue engineering: current state and prospects. Annu. Rev. Med. 52, 443–51 (2001).

4 Bruder S.P., et al. Mesenchymal stem cells in osteobiology and applied bone regeneration. ClinOrthop. Relat. Res. 355 Suppl, 247–56 (1998).

5 Ahmed T.A., Griffith M., Hincke M. Characterization and inhibition of fibrin hydrogel degrading enzymes during development of tissue engineering scaffolds. Tissue Eng. 13, 1469–77 (2007).

6 Ahmed T.A., Dare E.V., Hincke M. Fibrin: a versatile scaffold for tissue engineering applications. Tissue Eng. 14, 199–215 (2008).

7 Dare E.V., et al. Genipin cross-linked fibrin hydrogels for in vitro human articular cartilage tissue-engineered regeneration. Cells Tissues Organs. 190, 313–25 (2009).

8 Clines G.A. Prospects for osteoprogenitor stem cells in fracture repair and osteoporosis. Curr. Opin. Organ. Transplant. 15, 73–78 (2010).

9 Vilquin J.T., Rosset P. Mesenchymal stem cells in bone and cartilage repair: current status. Regen.Med. 1, 589–604 (2006).

10 Panetta N.J., Gupta D.M., Quarto N., Longaker M.T. Mesenchymal cells for skeletal tissue engineering. Panminerva Med. 51, 25–41 (2009).

11 Shrivats R.A., McDermott M.C., Hollinger J.O. Bone tissue engineering: state of the union. Drug.Discov. Today. 19, 781–6 (2014).

12 Stock U.A., Vacanti J.P. Tissue engineering: current state and prospects. Annu. Rev. Med. 52, 443–51 (2001).

13 Boo J.S., et al. Tissue-engineered bone using mesenchymal stem cells and a biodegradable scaffold. J. Craniofac. Surg. 13, 231–39 (2002).

14 Hao Z., et al. The scaffold microenvironment for stem cell based bone tissue engineering. Biomater.Sci. 5, 1382–92 (2017).

15 Hidaka S., et al. Royal jelly prevents osteoporosis in rats: beneficial effects in ovariectomy model and in bone tissue culture model. Evid. Based Complement. Alternat. Med. 3, 339–48 (2006).

16 Gasparotto V.P.O., et al. A new fibrin sealant as a three-dimensional scaffold candidate for mesenchymal stem cells. Stem Cell Res. Ther.5, 78–88 (2014).

17 Roseti L.,et al. Scaffolds for Bone Tissue Engineering: State of the art and new perspectives. Mater. Sci. Eng. C. Mater. Biol. Appl. 1, 1246–32 (2017).

18 Alving B.M., Weinstein M.J., Finlayson J.S., Menitove J.E., Fratantoni J.C. Fibrin sealant: summary of a conference on characteristics and clinical uses. Transfusion. 35, 783–90 (1995).

19 Spotnitz W.D. Fibrin Sealant: The Only Approved Hemostat, Sealant, and Adhesive-a Laboratory and Clinical Perspective. ISRN Surg. 203943; 10.1155/2014/203943 (2014).

20 Janmey P.A., Winer J.P., Weisel J.W. Fibrin gels and their clinical and bioengineering applications. J. R. Soc. Interface. 6, 1–10 (2009).

21 Barros L.C., et al. A new fibrin sealant from *Crotalus durissus terrificus* venom: applications in medicine. J. Toxicol. Environ. Health. 12, 553–71 (2009).

22 Thomazini-Santos I.A., Barraviera S.R.C.S., Mendes-Giannini M.J.S., Barraviera B. Surgicaladhesives. J Venom AnimToxins.7, 1–10 (2001).

23 Hino M., et al. Transmission of symptomatic parvovirus B19 infection by fibrin sealant used during surgery. Br. J. Haematol. 108, 194–5 (2000).

24 Kawamura M., Sawafuji M., Watanabe M., Horinouchi H., Kobayashi K. Frequency of transmission of human parvovirus B19 infection by fibrin sealant used during thoracic surgery. Ann. Thorac. Surg. 73, 1098–1100 (2002).

25 Dhillon S. Fibrin sealant (evicel(r) [quixil(r)/crosseal(tm)]): A review of its use as supportive treatment for haemostasis in surgery. Drugs. 71, 1893–1915 (2011).

26 Buchaim R. L. Effect of low-level laser therapy (LLLT) on peripheral nerve regeneration using fibrin glue derived from snake venom. Injury. 46, 655–60 (2015).

27 Cunha M.R.D., et al. Hydroxyapatite and a New fibrin sealant derived from snake venom as scaffold to treatment of cranial defects in rats. Mater. Res. 18, 196–203 (2015).

28 Machado E.G., et al. A new heterologous fibrin sealant as scaffold to recombinant human bone morphogenetic protein-2 (rhBMP-2) and natural latex proteins for the repair of tibial bone defects. ActaHistochem. 117, 288–96 (2015).

29 Buchaim D.V., et al. The new heterologous fibrin sealant in combination with low-level laser therapy (LLLT) in the repair of the buccal branch of the facial nerve. Lasers in Med. Science.25, 1–8 (2016).

30 Ferreira Jr R.S., et al. Heterologous fibrin sealant derived from snake venom: from bench to bedside – an overview. J. Venom. Anim. Toxins incl. Trop.23, 21; DOI10.1186/s40409-017-0109-8 (2017).

31 Biscola N.P., et al. Multiple uses of fibrin sealant for nervous system treatment following injury and disease. J. Venom. Anim. Toxins incl. Trop. (Online).23, 13; 10.1186/s40409-017-0103-1 (2017).

32 Cartarozzi L.P., et al. Mesenchymal stem cells engrafted in a fibrin scaffold stimulate Schwann cell reactivity and axonal regeneration following sciatic nerve tubulization. Brain Res. Bull. 112, 14–24, 2015.

33 Abbade L.P.F., et al. A new fibrin sealant derived from snake venom candidate to treat chronic venous ulcers. J. Am. Acad. Dermatol. 72, 5; https://doi.org/10.1016/j.jaad.2015.02.1081 (2015).

34 Orsi P.R., et al. A unique heterologous fibrin sealant (HFS) as a candidate biological scaffold for mesenchymal stem cells in osteoporotic rats. Stem Cell Res. &Ther. 8, 205; 10.1186/s13287-017-0654-7 (2017).

35 Ben-Ari A., et al. Isolation and implantation of bone marrow-derived mesenchymal stem cells with fibrin micro beads to repair a critical-size bone defect in mice. Tissue Eng. A. 15, 2537–46 (2009).

36 Langenbach F., et al. Improvement of the cell-loading efficiency of biomaterials by inoculation with stem cell-based microspheres, in osteogenesis. J. Biomater. Appl. 26, 549–64 (2012).

37 Khodakaram-Tafti A., Mehrabani D., and Shaterzadeh-Yazdi H. An overview on autologous fibrin glue in bone tissue engineering of maxillofacial surgery. Dent. Res. J. (Isfahan). 14, 79–86 (2017).

38 Roura S., Gálvez-Montón C., Bayes-Genis A. Fibrin, the preferred scaffold for cell transplantation after myocardial infarction? An old molecule with a new life. J. Tissue Eng. Regen. Med. 11, 2304–13 (2016).

39 Chang Y.S., et al. Mesenchymal stem cells for bronchopulmonary dysplasia: phase 1 dose-escalation clinical trial. J. Pediatr. 164, 966–72 (2014).

40 Lee J.W., et al. A randomized, open-label, multicenter trial for the safety and efficacy of adult mesenchymal stem cells after acute myocardial infarction. J. Korean Med. Sci. 29, 23–31 (2014).

41 Cooper J.A., Lu H.H., Ko F.K., Freeman J.W., Laurencin C.T. Fiber based tissue engineering scaffold for ligament replacement: design considerations and in vitro evaluation. Biomaterials.26, 1523–32 (2005).

42 Wei G., Ma P.X. Partially nanofibrous architecture of 3D tissue engineering scaffolds. Biomaterials. 30, 6426–34 (2009).

43 Yousefi A.M., Hoque M.E., Prasad R.G., Uth N. Current strategies in multiphasic scaffold design for osteochondral tissue engineering: a review. J. Biomed. Mater. Res. A. 103, 2460–81 (2014).

44 Tour G., Wendel M., and Tcacencu I. Cell-derived matrix enhances osteogenic properties of hydroxyapatite. Tissue Eng. Part A. 17, 127–37 (2010).

45 Dominici M., et al. Minimal criteria for defining multipotent mesenchymal stromal cells. The International Society for cellular Therapy position statement. Cytotherapy. 8, 315–7 (2006).

46 Junqueira L.C., Bignolas G., Brentani R.R. Picrosirius staining plus polarization microscopy, a specific method for collagen detection in tissue sections. Histochem. J. 11, 447–55 (1979).

47 Xu C., et al. A novel biomimetic composite scaffold hybridized with mesenchymal stem cells in repair of rat bone defects models. J. Biomed. Mat. Res. 95, 465–503 (2010).

48 Dayan D., Hiss Y., Hirshberg A., Bubis J.J., Wolman M. Are the polarization colors of picrosirius red-stained collagen determined only by the diameter of the fibers? Histochemistry. 93, 27–9 (1989).

49 Arakawa C., et al. Photopolymerizable chitosan-collagen hydrogels for bone tissue engineering. J. Tissue Eng. Regen. Med. 11, 164–74 (2017).

50 Laurencin C., Khan Y., EI-Amin S.F. Bone grafts substitutes. Expert Rev. Med. Devices. 3, 49–57 (2006).

51 Gholipourmalekabadi M., et al. In vitro and in vivo evaluations of three dimensional hydroxyapatite/silk fibroin nanocomposite scaffolds. Biotechnol. Appl. Biochem. 62, 441–50 (2015).

52 Kouhi M., Shamanian M., Fathi M., Samadikuchaksaraei Al., Mehdipour A. Synthesis, Characterization, In Vitro Bioactivity and Biocompatibility Evaluation of Hydroxyapatite/Bredigite (Ca7MgSi4O16) Composite Nanoparticles. J.O.M. 68, 1061–1070 (2016).

53 Barros L.C., et al. Biochemical and biological evaluation of gyroxin isolated from crotalus durissus terrificus venom. J. Venom Anim. Toxins Trop. Dis. 17, 23–33 (2011).

54 Spotnitz W.D., Prabhu R. Fibrin sealant tissue adhesive–review and update. J. Long Term. Eff. Med. Implants. 15, 245–70 (2005).

55 Spejo A. B., Neuroprotection and immunomodulation following intraspinal axotomy of motoneurons by treatment with adult mesenchymal stem cells. J. of Neuroinflammation. 15, 230 (2018).

56 Ahmed T.A., Giulivi A., Griffith M., Hincke M. Fibrin glues in combination with mesenchymal stem cells to develop a tissue-engineered cartilage substitute. Tissue Eng Part A. 17, 323–35 (2011).

57 Shapiro F. Bone developmented and its relation to factory repair. The role of mesenchymal osteoblasts and surface osteoblasts. Eur Cells and Mat. 2008;15:56–73.

58 Boxall SA, Jones E. Markers for characterization of bone marrow multipotential stromal cells. Stem Cells Int. 2012;2012:975871 [PMID:22666272 DOI:10.1155/2012/975871].

59 Casteilla L, Benard VP, Laharrague P, Cousin B. Adipose-derived stromal cells: their identity and uses in clinical trials, an update. World J Stem Cells. 2011;3(4):25–33.

60 Xiao Q, Wang SK, Tian H, Xin L, Zou ZG, Hu YL, et al. TNF-α increases bone marrow mesenchymal stem cell migration to ischemic tissues. Cell BiochemBiophys. 2012;62(3):409–14.

61 Mankani M.H., Kuznetsov S.A., Avila N.A., Kingman A., Robey P.G. Bone formation in transplants of human bone marrow stromal cells and hydroxyapatite-tricalcium phosphate: prediction with quantitative CT in mice. Radiology. 230, 369–76 (2004).

